# Improving antibody affinity using laboratory data with language model guided design

**DOI:** 10.1101/2023.09.13.557505

**Authors:** Ben Krause, Subu Subramanian, Tom Yuan, Marisa Yang, Aaron Sato, Nikhil Naik

**Affiliations:** Salesforce AI Research, Palo Alto, CA, USA; Twist Biopharma, Twist Bioscience, South San Francisco, CA, USA; Department of Biochemistry, Vanderbilt University School of Medicine, Nashville, TN, USA

## Abstract

Protein design involves navigating vast sequence spaces to discover sequences with desired traits. Language models (LMs) pretrained on universal protein datasets have shown potential to make this search space tractable. However, LMs trained solely on natural sequences have limitations in creating proteins with novel functions. In this work, we used a combination of methods to finetune pretrained LMs on laboratory data collected in an anti-CD40L single domain antibody library campaign to develop an ensemble scoring function to model the fitness landscape and guide the design of new antibodies. Laboratory experiments confirmed improved CD40L affinity in the designed antibodies. Notably, the designs improved the affinities of four antibodies, originally ranging from 1 nanomolar to 100 picomolar, all to below 25 picomolar, approaching the limit of detection. This work is a promising step towards realizing the potential of LMs to leverage laboratory data to develop improved treatments for diseases.

## 1 Introduction

Large scale language models (LMs) (Radford et al., 2018; 2019; Brown et al., 2020) initially developed for tackling challenges in natural language processing, have revolutionized many scientific disciplines by providing unprecedented capabilities for solving complex problems. Notably, ProGen (Madani et al., 2020) a language model trained on publicly available protein sequences, was able to successfully generate sequences of the bacteriolytic enzyme lysozyme that were very different in sequence space from any lysozyme in nature (Madani et al., 2023). More recently, Hie et al. (2023) harnessed pretrained language models to help guide antibody optimization. The search space for candidate proteins for any given application can be can be overwhelmingly vast, and these aforementioned successes show that language models have the ability to significantly reduce this search space.

While large language models trained on universal datasets can imitate the distribution of sequences in their training set, it is highly desirable to further train them to exceed the average quality of data in their training set. Furthermore, the distribution of sequences needed for downstream applications may significantly differ from what the model initially learned during pretraining. Applications of large scale language models in real world contexts have shown remarkable performance improvements upon fine-tuning the model using human feedback (Ziegler et al., 2019), wherein the model is further trained on high quality task specific responses or with reinforcement learning. For example, InstructGPT (Ouyang et al., 2022) used a combination of these two approaches to greatly improve language models responses and adaptability to make them useful for real world applications.

In the field of protein design, the closest analogy to learning from human feedback is learning from laboratory feedback, where the model is further trained to leverage laboratory measurements of desired properties. While pretraining teaches the language model to assign high probabilities to plausible protein sequences, the model remains agnostic to specific downstream tasks and struggles to generate proteins with novel functions absent in natural evolution—the training data. This limitation is especially significant for many pharmaceutical applications, where in many cases, no existing proteins in nature perform the desired function. Finetuning protein language models using laboratory data further refines these models to predict sequences that will perform the desired application with a higher level of functioning. Several methods have studied further training language models or using language model representations to finetune to laboratory data to better predict fitness (Hsu et al., 2022; Krause et al., 2021) or to design new proteins with higher fitness (Biswas et al., 2021), with promising results. However, finetuning language models using laboratory data in systems that have a direct clinical application remains an under-explored area.

Antibodies are proteins produced by the immune system to defend against foreign agents. The immune system can, in principle, generate antibody variants to bind any target antigen with high affinity and specificity, which also makes antibodies ideal candidates for therapeutic applications (Chiu et al., 2019). There are currently over 100 FDA-approved antibodies that are used to treat a wide range of diseases and are often the best-in-class therapeutic agent (Mullard, 2021). Despite their utility in the clinic, the process of lead optimization going from a candidate antibody to variants that have desired properties is laborious and time consuming. This is because, like in other protein engineering cases, the search space is vast. Combining language models trained on protein sequences with experimentally determined can potentially accelerate the antibody design process. Recent work has shown promise in designing antibodies using deep learning and language model based methods (Bachas et al., 2022; Porebski et al., 2023; Shanehsazzadeh et al., 2023). Here, we finetune a protein language model with laboratory data from a library campaign to design anti-CD40L single domain antibodies. We design multiple scoring functions using a combination of finetuned pretrained language models and other models that condition on language modeling feature representations. We use these scoring functions to guide the design of new CD40L binding antibodies, and validate our methods with laboratory results that show that we significantly improve the binding affinity of several of the original hits that were already strong binders. We describe our high level approach in Figure 1.

**Figure 1:**
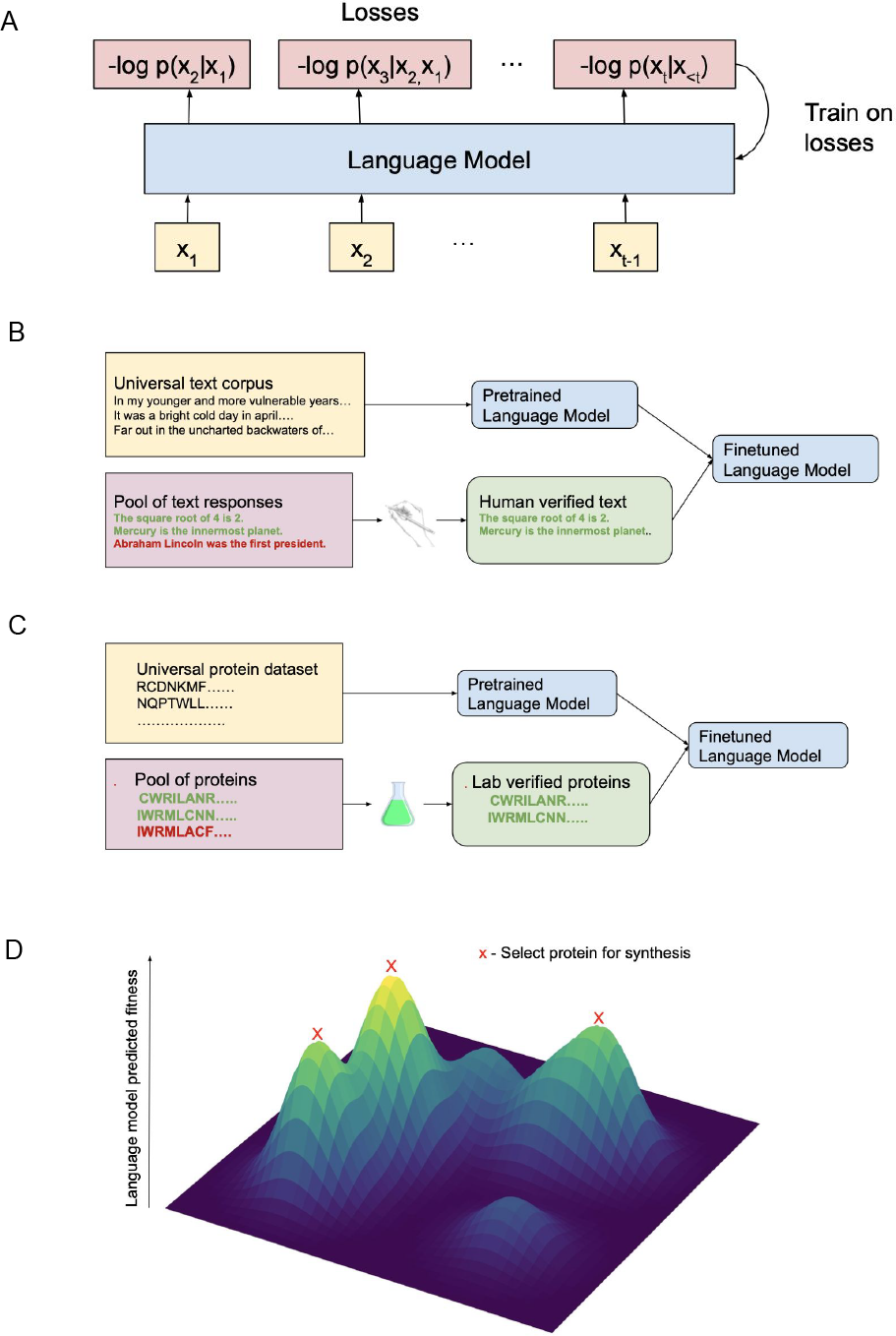
(A) Language models process the elements of a sequence one-by-one, and iteratively predict the probability distribution over next element *x*_*t*_ conditioned on the previous elements *x*_*<t*_. They are trained to minimize the the negative log-likelihood loss of sequences in their training set using the gradient of the loss with respect to the model parameters. (B) Textual language models learn a universal representation of human generated text from their pretraining phase. However, they are much more useful for applications when they are also finetuned to high quality human verified text that is relevant for the types of tasks they will later be asked to perform. (C) Pretrained protein language models learn a universal representation of proteins, allowing them to distinguish between plausible and implausible protein sequences. Further refinement on laboratory verified sequences that perform particular functions should help them learn the distribution of plausible proteins that perform a particular function at a high level (D) Predicted fitness landscapes can be used to guide selection of candidate proteins. In this work, we used laboratory data to finetune language models to predict a fitness landscape, and found a diverse set of candidate proteins with high predicted fitness that were synthesized and tested in the laboratory.

## 2 Results

Cluster of differentiation 40 (CD40) is a cell surface protein primarily found on antigen-presenting cells such as B cells, dendritic cells, and macrophages. Its ligand, CD40L (also known as CD154), is predominantly expressed on activated T cells. The interaction between CD40 and CD40L coordinates and regulates key immune responses to various challenges, including infections and autoimmune disorders (Fig. 2A). Crystallographic views of the CD40-CD40L complex show a 2:3 stoichiometry for the interaction (An et al., 2011)(Fig. 2B). For patients suffering from autoimmune disorders, disrupting the interaction between CD40-CD40L using antibodies is potentially a powerful therapeutic intervention (Daoussis et al., 2004; Law & Grewal, 2009).

**Figure 2:**
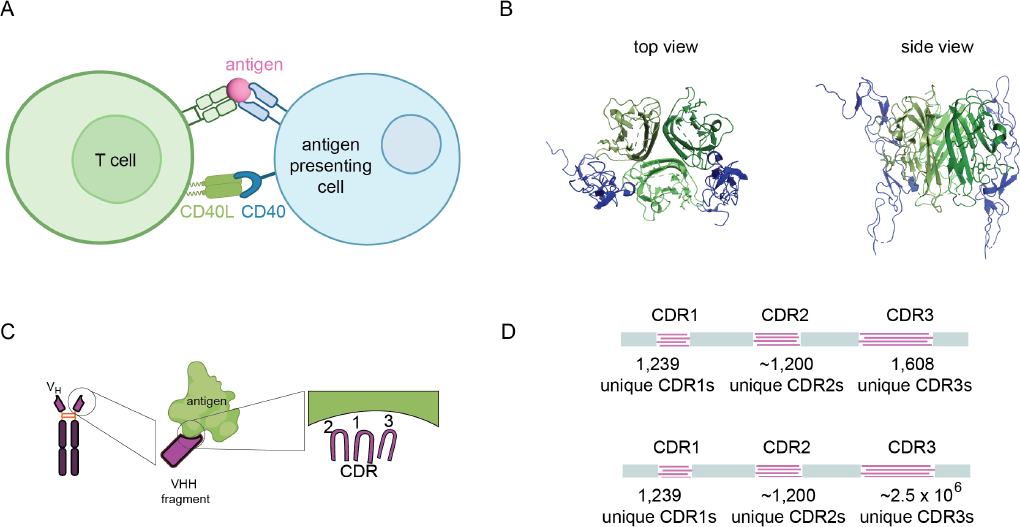
Screening nanobody binders to CD40L. (A) Schematic depicting the interaction between CD40 expressed on the surface of an antigen presenting cell and the CD40L present on the surface of a T cell, which brings the T-cell and antigen presenting cell together to facilitate an immune response. (B) Crystallographic views (from top and side) of the CD40-CD40L complex,, with CD40 in blue and CD40L in green. Based on PDB ID (3QD6) (An et al., 2011). (C) Nanobody schematic, showing the naturally occurring camelid antibody, the VHH fragment that binds antigens, and the three CDR loops of the VHH fragment that interacts with the antigen. (D) Schematic of the two antibody library sequences used in the study. The unique CDRs are combinatorially joined to create a large set of antibodies for screening.

**Figure 3:**
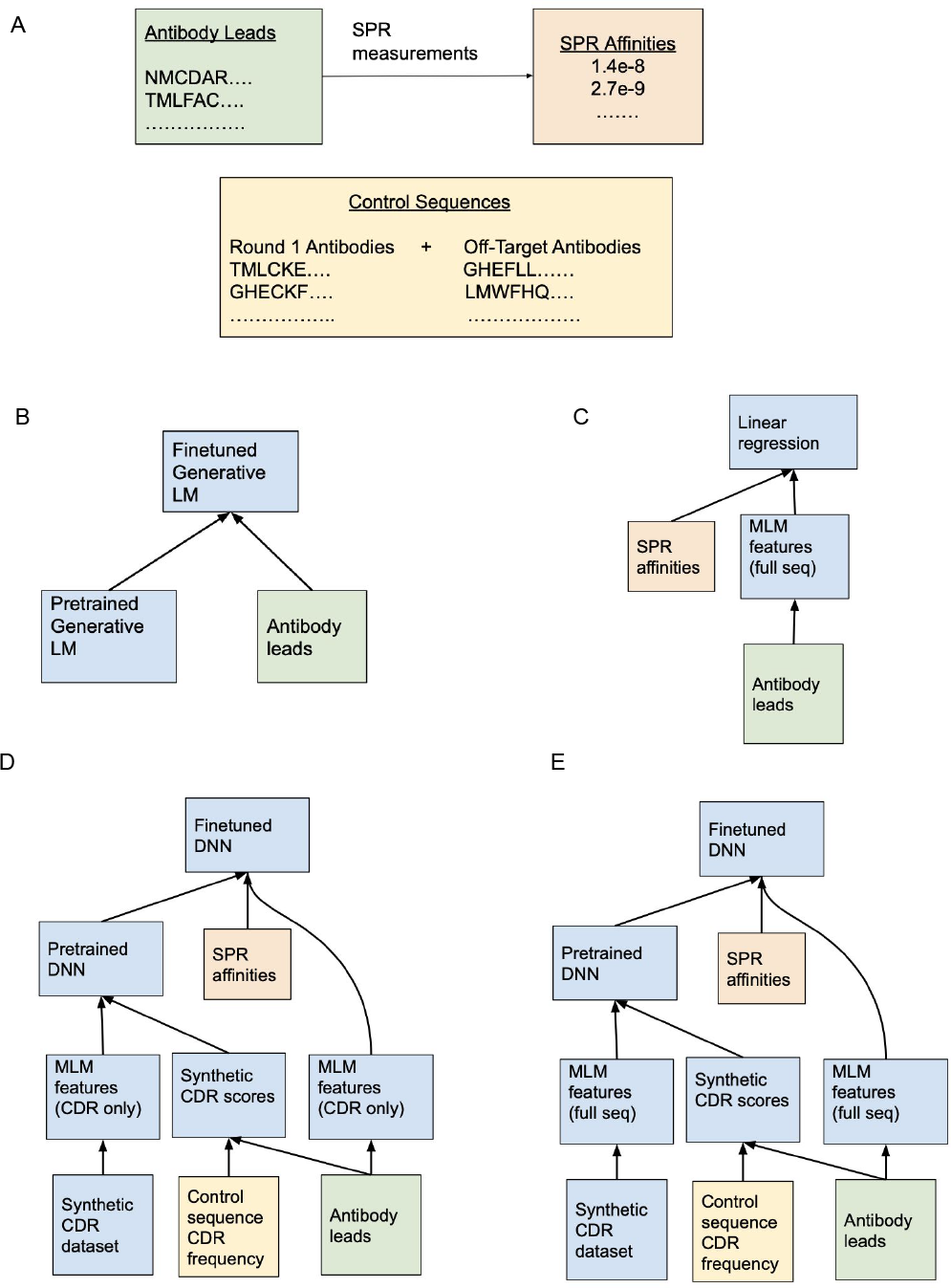
(A) Data sources included the top antibody leads found from NGS selection and their corresponding binding affinities, as well as “control sequences” present in the first round of selection and the final round of 4 off-target antibody campaigns. The control sequences are more likely to be over-represented in the library or have non-specific binding. (B-E) We used an ensemble of 4 scoring models to rank candidate antibodies. These graphs depict the data used to train each of these models. (B) Data pipeline for CDR Generative LM scores. We finetuned a pretrained language model to a CDR only representation the top antibody leads found in the laboratory campaign. The log likelihoods output by the language model were used as a scoring function. (C) Data pipeline for VHH-Feature regression scores. We applied linear ridge regression using a pretrained mask language model to compute a feature representation of full antibody sequences, and using normalized SPR affinities as labels for prediction. (D) Data pipeline for CDR-feature NGS-pretrained DNN scores. We developed a synthetic scoring system based on the frequency of CDR sequences in the antibody leads divided by the frequency of the same CDR sequences in the control sequences. We pretrained a DNN to predict these synthetic scores and finetuned them to predict binding affinity of the antibody leads from their CDR sequences. (E) Data pipeline for VHH-feature NGS-pretrained DNN scores. We applied similar methods to (D) but with the full VHH sequences.

**Figure 4:**
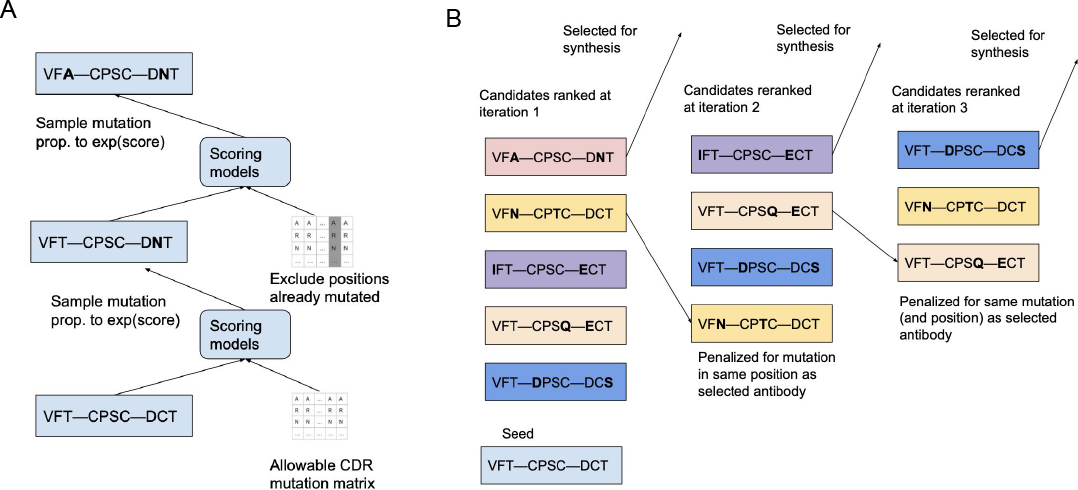
Designing and selecting mutational trajectories. (A) We designed candidate mutational trajectories from seed sequences by first considering and scoring the set of all possible single mutations in CDRs. We then sampled a mutation with a probability proportional to *e* to the power of the mutations score. After sampling the first mutation, we sampled up to 4 more mutations in the same fashion, but excluding positions that had already been mutated in previous iterations. (B) After generating a large set of candidate mutational trajectories, we selected sequences to be synthesized for laboratory testing. At each selection iteration, we selected the top scoring sequence, and added a penalty to the scores of sequences that had the same mutation(s) as the selected sequences, and mutations at the same position(s) as the selected sequence. This ensured diversity in our selected set. We performed 15 selection iterations for each of our 5 seed antibodies.

In most animals, antibodies bind their target with high specificity using the variable light chain and the variable heavy chain. The antibodies in camelids are an exception in that these antibodies do not possess a light chain, using only the variable heavy chain (termed the VHH fragment) to bind the target antigen with high specificity (Hamers-Casterman et al., 1993). Over the last decade, the VHH fragment (*∼*120 amino acid residues) have become more widely studied due to desirable properties, including high solubility, their overall ease of synthesis and access to dense tissues or structures due to their small size (Könning et al., 2017; Smolarek et al., 2012; Chanier & Chames, 2019). The VHH fragments are also known as single-domain antibodies (sdAb) or nanobodies (Fig. 2C). Recently, large libraries of nanobody sequences displayed on bacteriophages have been developed for use as a starting library to screen binders to target antigens (Romao et al., 2016; Yuan et al., 2022; Moreno et al., 2022). Our designs and training data consisted entirely of VHH fragments, which we refer to as nanobodies and antibodies interchangeably throughout the paper.

We began by screening two phage displayed nanobody libraries for CD40L binding generated by Twist Bioscience (Yuan et al., 2022). Both libraries were constructed on a partially humanized VHH framework to lower immunogenecity for therapeutic development (Ter Meulen et al., 2006). One library shuffled *∼*10^3^ llama-based variants for each of the three complementarity-determining regions (CDRs), resulting in a library with *∼*3*×*10^9^ unique nanobody sequences (Yuan et al., 2022). The other library used *∼*10^3^ llama-based variants for CDR1 and CDR2, and *∼*2.5*×*10^6^ unique human CDR3 variants identified from naive and memory B-cells and yielded a library of *>*10^10^ unique variants (Yuan et al., 2022). The two libraries were pooled together in an antibody discovery campaign to identify CD40L binders. The data generated from the campaign comprised:

1. NGS selection rounds 1-4 for CD40L antibodies. These selection rounds use phage display (Smith & Petrenko, 1997) to encode the protein in the phage’s genome in a location that will cause it to appear on the surface of a phage. At each selection round, the phages that encode for antibodies are passed over the target (which is CD40L protein), and the phages that stick are sequenced using next generation sequencing (NGS). This allows on the order of billions of proteins to be screened at once. At each round of selection, stricter conditions are used (more washes, less target). As a result, phages that encode for proteins that are tighter binders will be relatively more frequent in the later rounds of selection. 2. A set of 60 lead antibodies were identified as top hits through a combination of NGS selection and subsequent ELISA screenings. Surface plasmon resonance (SPR) was used to measure their binding affinity to CD40L, indicated by the dissociation constant (*K*_*D*_). A lower *K*_*D*_ signifies tighter binding. 3. NGS selection rounds 1-4 for 4 other antibody targets that used the same library design. Antibodies that bind to many targets may be binding non-selectively.

Comprehensive details of our laboratory data collection are available in Section 4.3. Our objective was to use this data to train a scoring system to predict antibody binding strength to the target, and then use that scoring system to guide the design of novel antibodies with improved affinity.

We observed that the likelihoods predicted by an unmodified pretrained language model were not well correlated with the affinity of lead antibodies, motivating us to finetune models using the laboratory data. To do this, we trained 4 distinct scoring models that each used differing information. Since we only had the opportunity for 1 round of synthesis and laboratory testing, we wanted to ensure that our designs were robust to flaws in any individual scoring model. We also hoped to that using a combination of scoring models would yield information about which model was the most useful for improving affinity.

The first scoring model used a causal language model (LM) to predict the likelihood of sequences. Causal language models can model a protein of length *T* as a string of amino acids *x* = [*x*_1_,.., *x*_*T*_], where the likelihood of a protein *x* under the model is given by

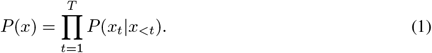

LMs can be trained to maximize the expectation of *log*(*P* (*x*)) using samples of *x* that come from the training set. A well trained LM will assign a higher probability to sequences that have similar patterns to the sequences that occur in the training set. We leveraged the pretrained Transformer (Vaswani et al., 2017) language model ProGen (Madani et al., 2020), which should assign a high probability to proteins with patterns that match what is found in nature because it is trained on large datasets of natural proteins.

We found that ProGen’s likelihoods correlated very weakly and insignificantly with binding affinity of our 60 antibody leads (Spearman *r* = 0.08, *p* = 0.5), suggesting that pretrained models without modification may not be strong predictors of fitness in this particular setting. To improve ProGen’s ability to recognize patterns specific to CD40L antibodies, we finetuned it to the sequences of the lead CD40L antibodies from our laboratory data. When modeling these sequences with ProGen, we used only the CDRs to represent the sequence, omitting the remainder (framework) of the variable heavy chain sequence. The CDRs of an antibody typically interact with the antigen, and are therefore especially important for predicting binding affinity. The log-likelihoods predicted by our finetuned LM, which we refer to as CDR generative LM scores, were the first of four scores that we used to rank possible antibody candidates.

Our remaining three models used representations from a pretrained masked LM (Devlin et al., 2018), ESM-1v (Meier et al., 2021). Masked LMs are trained to predict masked amino acids in protein sequences in bi-directional fashion, and can learn useful vector representations of proteins. Our second score applied ESM-1v to the full heavy chain sequence of the 60 lead antibodies, mapping each sequence of amino acids to a sequence of vectors. We applied mean pooling across the sequence to these vectors to obtain a single vector. We applied linear ridge regression to predict the binding affinities of the lead antibodies from the vector representations of the sequences. We refer to the predictions of this model as VHH-feature regression scores, and used it as the second of 4 scores to rank antibody candidates.

Our third score used information from the NGS selection campaigns as well as from the final antibody leads. We developed a pseudo data generation and corresponding pseudo labeling technique that correlated with our SPR binding affinity data that we later used to help train models to predict binding affinity. We specifically considered the set of every possible combination of CDR sequences found in the antibody leads. Some CDR sequences seemed to predict higher affinity than others. We developed a pseudo-score for an antibody based on its CDRs given by the ratio of how often each CDR sequence was present in the antibody leads (indicating higher fitness) divided by how often it was present in round 1 of the CD40L campaign and the final round of the other 4 library campaigns (likely indicating over-representation of the library or non-specific binding). This ratio was then used to assign a score to random new combinations of CDR sequences to create a synthetic training set that used information from the NGS selection. We trained a deep neural network (DNN) to rank the relative fitness of the synthetic data with the pseudo-scores, and then finetuned it to predict the ranking of the SPR-measured affinity of the antibody leads. We used a CDR only representation of the antibodies for this model. Like with the feature regression scores, we also used vector representations (with mean-pooling) computed with ESM-1v as inputs to the model. We refer to this as CDR-feature NGS-pretrained DNN scores.

Our fourth score used the same strategy as the third score, but trained the model on full variable heavy chain sequences. For generating synthetic data, we used the randomly recombined CDR sequences and slotted them in the variable heavy chain sequences of random antibody leads from our set of 60. Pretraining on full sequences more naturally allowed the model to be finetuned to the full length antibody leads. We refer to this score as VHH-feature NGS-pretrained DNN scores.

Our final score used a weighted combination of our 4 scoring models, which we refer to as our ensemble score. Our final weighted scores leveraged the CDR generative LM scores most heavily (see Table 2). We used two different strategies for using the ensemble scores to guide the design of new antibodies. Our first strategy used the score to make a series of mutations to previous antibody leads, and our second strategy used the score to design antibodies with novel combinations of the CDR sequences found in the antibody leads. We designed 96 antibodies in total for laboratory testing; 75 using mutations from antibody leads and 21 from recombining CDR sequences.

To generate antibodies using series of mutations from antibody leads, we generated mutational trajectories by iteratively sampling mutations in CDRs one at a time using an energy function based on our ensemble scores. At each iteration, we computed the ensemble score of all the sequences that could result from adding a single mutation to the sequence at the previous iteration, excluding positions where a mutation from the seed sequence had already occurred in a previous iteration to ensure *n* iterations gives *n* mutations. We sampled 256 sequences each of 1-5 mutations from 5 different seed antibodies and used this as a candidate pool for sequences to be synthesised and tested. We selected 3/256 sequences from each setting (1,2,3,4 and 5 mutations from 5 different antibodies) to synthesize, resulting in 15 per seed sequence and 75 total. For each seed sequence, we selected the 15 sequences for synthesis one-by-one. We initialized scores for each candidate antibody to be equal to its corresponding ensemble score. At each selection iteration, we selected the antibody with the highest score. Then we subtracted penalties to the scores for the remaining antibodies that had 1. the same mutation(s) as the selected antibody 2. mutations in the same position(s) as the selected antibody To ensure that we selected exactly 3 of each number of mutations, we also removed all n-mutation antibodies from the pool once 3 n-mutation antibodies had been selected. We repeated this process for the candidates from each of the 5 seed sequences.

To generate antibodies by re-combining CDR sequences, we considered a set of all the CDR sequences in the 3 regions present in the antibody leads. We then applied all permutations of those CDR sequences to create a large set of candidate antibodies. We computed scores for all of these candidates based on our CDR generative language model and our CDR-feature NGS-pretrained DNN (we excluded the full sequence models since we only considered CDR only representations at this phase). To select the final antibodies, we iteratively selected the highest scoring CDR sequence combinations, and updated the scores at each iteration to penalize sequences that contained one or more of the same CDR sequences as the selected sequences. We selected 21 sequences in total through these iterations. Since we had been working with a CDR only representation of the design, we needed to fill in the remainder of the variable heavy chain sequences. To do this, for each of the 21 CDR-only sequences, we found the sequence in the antibody leads with the most overlapping CDR sequences. We used the SPR affinity of the leads to break ties when more than one antibody lead sequence tied for the most overlap with the designed sequence. We then designed the final sequences using the CDR sequences from the selected sequences combined with the remainder of the variable heavy chain sequence from the chosen antibody leads.

We tested our 96 designed antibodies for affinity to CD40L using SPR. SPR measures the dissociation constant (*K*_*D*_) of 2 binders *A* and *B*. The *K*_*D*_ value for *A* and *B* depends on the equilibrium concentrations of *A* and *B* (given by [*A*] and [*B*]) and the concentration of A and B bound together (given by [*AB*]), and is computed using the ratio

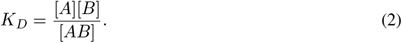

When using SPR, *K*_*D*_ is calculated using the ratio *K*_*off*_ */K*_*on*_, where *K*_*on*_ is the rate at which *A* binds to *B*, and *K*_*off*_ is the rate at which bound *A* and *B* dissociates. Our experiments had a limit of detection for *K*_*off*_ of 10^*−*5^, meaning that any antibody with a *K*_*off*_ of less than or equal to 10^*−*5^ uses *K*_*off*_ = 10^*−*5^ for *K*_*D*_ calculations. In these cases where an antibody reaches the limit of detection for *K*_*off*_, the actual binding affinity could be stronger than what is indicated by the measured *K*_*D*_ value. There were 4 such antibodies in our original set of leads, including 1 of the seed antibodies used for generation.

We were able to obtain *K*_*D*_ values for 83*/*96 tested antibodies. 54*/*96 of designed antibodies were shown to have tighter than nanomolar affinity (*K*_*D*_ *<* 10^*−*9^), as compared with 21*/*60 of the original antibody leads. 16 of the designed antibodies were at the limit of detection for *K*_*off*_, suggesting that they could have a higher affinity than indicated by their calculated *K*_*D*_ values. A plot of the affinities of the set of antibodies designed by applying mutational trajectories to the 5 seed sequences is given in Figure 5C. The designs were able to improve affinity the seed sequences by up to 40x. The resulting antibodies derived from all 5 seed sequences included antibodies that were less than 25 picomolar, and were at the limit of detection for *K*_*off*_. The only antibody that was not improved started at the limit of detection for *K*_*off*_. While the overall success rate of the CDR recombination antibodies was lower (Figure 5A), there were still 2 high affinity hits at the limit of detection for *K*_*off*_, and below nanomolar affinity binders that were up to 8 mutations away from the nearest antibody lead in the training set (Figure 5B). We also analyzed the performance of the 4 scoring modules in predicting the affinity of designed sequences in Table 1, and found that the finetuned CDR generative LM was the only scoring system that correlated with binding affinity within the set of designed sequences. Since the designed sequences are selected to have high scores, it is expected that any correlations between binding affinity of designed sequences and scores would be reduced as compared with sequences that are designed independently of the scores.

**Table 1:** Spearman correlation of 4 scoring models with binding affinity on held-out antibody leads from library, and generated antibody sequences. The NGS-pretrained models scored the highest on predicting the affinities of lead antibodies. Since the generated sequences are selected to have high scores, it is expected that any correlations between binding affinity of designed sequences and scores would be reduced (as compared with sequences that are designed independently of the scores). We find that even so, The CDR language modeling likelihoods still correlated significantly with affinity in this setting (*p <* 0.01), suggesting that it may have been the most important scoring component in driving affinity improvements.

| Model | Antibody lead Spearman | Generated antibody Spearman |
| --- | --- | --- |
| CDR generative LM | 0.35 | 0.33 |
| CDR-feature NGS-pretrained DNN | 0.54 | -0.10 |
| VHH-feature NGS-pretrained DNN | 0.52 | 0.02 |
| VHH-feature linear regression | 0.44 | 0.02 |

**Figure 5:**
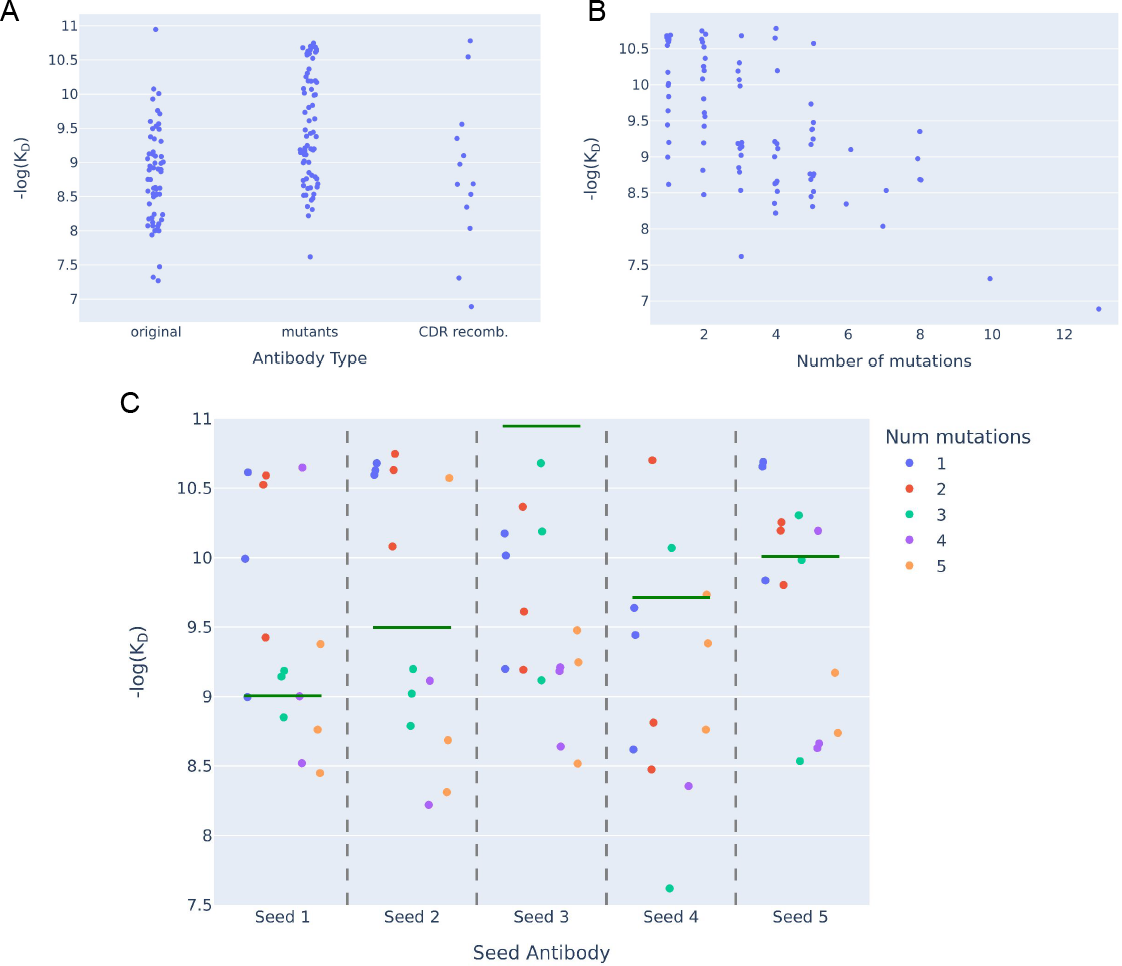
SPR binding affinities of generated proteins. (A) SPR binding affinity of original set of antibody leads vs. mutational trajectories of seed antibodies vs. antibodies generated via CDR recombination. In general, mutations of antibody leads had higher affinity, but CDR recombination antibodies had more novel hits. (B) SPR binding affinity plotted against edit distance to nearest antibody leads. The generated antibodies included strong binders as many as 8 mutations away from the nearest pre-existing antibody in the training set. (C) SPR binding affinity of antibodies (dots on graph) vs. their respective seed sequences (horizontal lines on graph). Our models were able to improve the binding affinity of 4/5 seed antibodies, by a factor of up to 40 fold.

## 3 Discussion

We used a combination of pretrained language models and laboratory data to design new CD40L binding single-domain antibodies. We initially found that an unmodified pretrained language model was a poor predictor of fitness of antibodies in our training set, thus supporting the motivation of finetuning models using data from the initial CD40L library. We developed an ensemble of 4 scoring functions, 1 based on finetuning a pretrained language model directly and 3 based on training models that condition on features from pretrained language models, to model the fitness landscape of CD40L binders. We implemented a strategy to design a diverse set of candidates with high predicted fitness according to our ensemble models. Results show that the designed antibodies were able to improve affinity of the seed antibodies that we used as starting points by up to 40 fold, and that using the ensemble score to design novel CDR re-combinations yielded sub nanomolar affinity hits as many as 8 mutations away from the nearest sequence in our training set. These results indicate that the ensemble score was able to model the fitness landscape well enough to be useful for designing high affinity sequences. Further analysis showed that the CDR generative LM scores were correlated with binding affinity on our designed sequences, whereas the 3 scores that used models trained to condition on language model representations were not. This suggests that the CDR LM scores the most important in driving the observed improvements to affinity, and that finetuning a pretrained language model directly was more useful than training models to condition on language model features in this setting. The results suggest training language models to predict the fitness landscape and using that predicted landscape to guide design is an effective way for language models to leverage laboratory data.

Our models in this work did not directly use the target (CD40L) sequence or structure, or explicitly use the structure of the antibody sequence. Methods that are agnostic to this information are especially useful for settings where the target structure or binding is not known. However, for problems where the structure of the binding pocket of the target is known or the complex between the target and candidate antibody can be accurately predicted, models that can also leverage structure explicitly could have an advantage in predicting strong binders. Recent advances in predicting the structure of proteins (Jumper et al., 2021; Baek et al., 2021; Lin et al., 2022) and protein complexes (Evans et al., 2021) could be useful in models that predict whether an antibody will bind to a target. Information output from these models could be used to create additional scoring functions that predict how well antibodies would be expected to bind. Another approach is to use generative models that condition directly on structural information to model likelihoods of protein sequences (Hsu et al., 2022; Dauparas et al., 2022). Diffusion models (Ho et al., 2020) can be used to generate partial structures for protein-protein complexes (Watson et al., 2023), which can then be conditioned on using LMs or other generative models to model sequence likelihoods. These approaches are promising for developing initial candidates to solve difficult design problems that involve protein-protein interactions, however finetuning from laboratory data will likely be important for applying these approaches to develop optimized therapeutics that are useful in practice.

Our approach could also be used for machine learning guided directed evolution. Directed evolution (Arnold, 1998) is a protein engineering method that iterates between selecting the most fit proteins and mutating them to design more candidates. Machine learning based approaches can be used to help guide the selection candidates to make this process more efficient (Yang et al., 2019; Ma et al., 2021). Like our work in this paper, some previous works have focused on single shot data driven design, with the goal of generating the highest fitness proteins possible from a single round of data (Fannjiang & Listgarten, 2020; Chan et al., 2021).

In conclusion, we demonstrated a framework for language models to leverage laboratory data to design single-domain antibodies that have potential real world applications towards treating diseases. Designing therapeutics in practice often requires designing proteins that satisfy many criteria, including binding to the target as well as maximizing stability while minimizing immunogenicity. Our methods demonstrated the capability to increase binding affinity to the target, which could directly improve the effectiveness of existing therapeutics. Our approach was also able to increase the number and diversity of hits that bind to the target, which could make it easier to find candidates that satisfy other important criteria.

## 4 Methods

We leveraged data from a CD40L library campaign to design new anti-CD40L antibodies. Our strategy for designing new antibodies was to:

1. train several different models that are able to predict SPR binding affinity on the top hits from the original library and combine them to create a scoring system.
2. sample a large set of sequences within a constrained space that score highly under this scoring system. For our constrained space, we consider A. antibodies that are a specific number of mutations away from top hits in the original library and B. antibodies that use combinations of CDR sequences present in the original library.
3. select top scoring sequences for synthesis one-by-one out of the larger set, while adding a penalty to ensure that each selected sequence is different enough from each previously selected sequence.

### 4.1 Scoring system

For our scoring system, we aimed to use a combination of models that use different inductive biases but all perform well at predicting the SPR binding affinity of the antibody leads, as measured by as measured by *K*_*D*_ values. We considered approaches that leveraged the following data sources:

1. The measured affinity values of the 60 antibody leads. This is the most direct signal that directly indicates binding affinity. We denote this dataset as *D*_*a*_ = [(*x*^(1)^, *y*^(1)^), …, (*x*^(60)^, *y*^(60)^)]
2. The presence or absence of an antibody in the final set of leads. Antibodies that appear in the leads tend to be much stronger binders–so we would like to identify patterns in common between these antibodies to predict additional strong binders. This is simply the unlabeled sequences from *D*_*a*_ given by [*x*^(1)^,.., *x*^(60)^].
3. The frequency of a CDR sequence appearing in the final set of leads. CDR sequences that occur frequently in the final leads would be thought to help make binding stronger. Since there are 3 different CDRs in each antibody, we denote 3 sets, 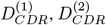, and 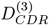, where 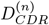 is a dataset of pairs of CDR sequences *s* and their frequencies *f* in the *n*th CDR, given by 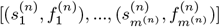, where *m*^(*n*)^ gives the number of unique sequences in the *n*th CDR.
4. The Frequency of a CDR sequence appearing in control sequences, comprising of round 1 of biopanning and the final round of off-target campaigns. Spans that occur frequently in round 1 may be over-represented in the library, and spans that are present in the final round of other binding campaigns may indicate non-specific binding. We counted the total number of unique sequences containing each CDR sequence in the first round of biopanning and in round 4 of 4 different off-target binding campaigns. We summed these frequencies for each span to create datasets of CDR sequences and corresponding frequencies. Like in the previous paragraph for lead frequencies, we denote 3 sets, 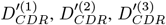, where 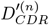 is a dataset of pairs of CDR sequences *s* and their frequencies *f* ^*′*^ in the *n*th CDR given by 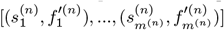. Note that the 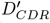 sets contain the same CDR sequences *s* as in the *D*_*CDR*_ sets, but have different frequencies paired with each CDR sequence.

We trained several different models using these datasets and used the ones that predicted SPR well scores empirically. We developed 4 scores, which we refer to as CDR Generative LM scores, VHH-feature regression scores, CDR-feature NGS-pretrained DNN scores, and VHH-feature NGS-pretrained DNN scores, that we combined and used to guide antibody selection.

#### 4.1.1 CDR Generative LM scores

The final 60 antibody leads correspond to the winners of the library campaign. It is desirable for new designs to contain similar patterns to this set of winners. Motivated by this insight, we finetuned a generative LM (ProGen) to the concatenated CDR sequences of the final leads. ProGen is pretrained on large databases of existing proteins found in nature, which should by default cause it to assign a higher probability to more plausible proteins than less plausible ones, as well as give it a useful feature representation for the space of proteins. Generative language models predict a conditional distribution over the next amino acid *x*_*t*_ conditioned on the previous amino acids *x*_*<t*_, given by *P* (*x*_*t*_ *x*_*<t*_). They can assign a score to a assign scores to proteins given by the log probability of that sequence under the language model, given by

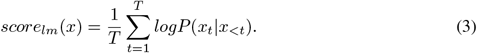

We finetuned ProGen to the concatenated CDR sequences of the antibody leads that included the framework of the variable heavy chain sequences. This finetuning minimized a loss function given by *score*_*lm*_, thus minimizing the negative log likelihood and maximizing the scores of the CDR-only antibody leads. Finetuning used the Adam optimizer (Kingma & Ba, 2014) with learning rate 10^*−*4^ and gradient norm clipping (Pascanu et al., 2013) with a norm threshold of 0.25. We found that the antibody lead-finetuned version of ProGen achieved a Spearman correlation of 0.35 with the SPR affinities, despite not being trained directly on any affinity labeled data.

#### 4.1.2 VHH-feature regression scores

This score used linear ridge regression on top of a feature representation of the variable heavy chain sequences. We used the ESM-1v (Meier et al., 2021) pretrained mask language model to map each sequence to a feature vector. We then fit linear ridge regression to predict normalized *K*_*D*_ values from the feature vector. The ridge parameter was tuned using cross validation. The scores were given by:

1. Feed *x* into the ESM-1v masked language model. This maps the sequence of amino acids in *x*, given by *x*_1_,.., *x*_*T*_, (where *T* is the number of amino acids in *x*), to a sequence of vectors *v*_1_,.., *v*_*T*_. We apply mean pooling across the sequence to give *v* = *mean*(*v*_1_,.., *v*_*T*_), giving a vector representation of the sequence.
2. Apply inner product between the learned weights and the vector representation of the sequence, given by *w*^*T*^ *v*, to give the final feature regression score for sequence *x*.

When applying 8 fold cross validation with training set sizes of 32 and validation set sizes of 28, this method achieved an average Spearman correlation of 0.44 with SPR affinity of the held out examples. The final linear weights *w* were fit using all 60 supervised training examples with the ridge parameter found from cross validation.

#### 4.1.3 CDR-feature NGS-pretrained DNN scores

Our next score jointly leveraged information from the NGS reads from the biopanning campaign along with information from the SPR affinity measurements on the leads. For this score, we converted each sequence *x* to a concatenation of the CDR sequences *x*_*CDR*_. This representation only includes amino acids in CDRs–which are the most important for determining affinity to targets–and excludes amino acids in non-CDRs of the variable heavy chain sequence. We mapped each *x*_*CDR*_ to a vector *v*_*CDR*_ by passing each sequence to a feature representation with ESM-1V and applying mean pooling as we did to compute the linear regression scores. We then created a synthetic pretraining dataset based on pseudo labels from the CDR frequencies to pretrain a deep neural network (DNN) to predict fitness. To do this, we recombined CDR sequences from the *D*_*CDR*_ dataset to create new concatenated CDR sequences. Each synthetic concatenated CDR sequence *x*_*synth*_ was generated using *concatenate* 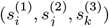 where *i,j*, and *k* are randomly sampled from the set of possible indices. The pseudo score *y*_*synth*_ for sequence *x*_*synth*_ is then given by

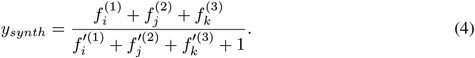

This ratio favors sequences with CDRs that are frequently present in the leads, but not overrepresented in the initial library or frequently binding to other targets. We found that when applied to the antibody leads, the numerator in Equation 4 correlated with binding affinity with a Spearman of *r* = 0.32, whereas including the full ratio improved it to 0.47. Note that Equation 4 alone can only be used to score sequences that only consist of CDRs present in the set of leads, and cannot generalize to the majority of the possible sequence design space.

We pretrain a simple deep neural network (DNN) with 2 hidden layers to predict *y*_*synth*_ from *x*_*synth*_ (where *x* synth is mapped to a vector using ESM-1v as with the feature regression scores before being input to the DNN). Our goals were to 1. Interpolate between the pseudo scores of CDRs in a way that can generalize to new sequences that contain CDRs not present in the leads 2. learn a useful pretrained representation that can later be finetuned to SPR binding affinity that accounts for information from the frequency of CDRs in the initial and final rounds of the CD40L campaign as well as the final round of off-target campaigns.

We care more about correctly predicting the relative ranking of fitness than predicting absolute scores, and we would also like to later finetune the model to SPR *K*_*D*_ values that are on a completely different scale. Therefore, we used a Bradley-Terry (Bradley & Terry, 1952) based cost function to train the DNN to correctly classify the relative ranking of fitness of pairs of synthetic samples. Bradley-Terry based cost functions are often used to train reward models on preferences (Christiano et al., 2017).

At each training iteration, we sample 2 training examples from the synthetic dataset. We denote the sequence with the higher fitness as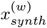, and the sequence with the lower fitness as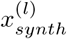. After applying ESM-1v and mean pooling to 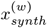 and 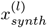, the DNN is used to map the resulting vectors to a single score value. The loss given by this pair is given by

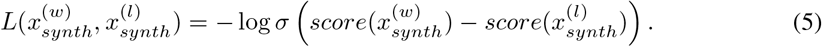

We trained the DNN on a dataset of 9000 of these synthetic examples using the above cost function and the Adam optimizer (Kingma & Ba, 2014) with learning rate 10^*−*5^ and 100 epochs. We then finetuned this pretrained DNN directly on our SPR binding affinity data. For finetuning, we only sampled *x* from the training set given by the SPR data, and use the ranking of the true SPR value *y* (as opposed to *y*_*synth*_ to determine *x*^(*w*)^ and *x*^(*l*)^. The cost function for finetuning was therefore given by

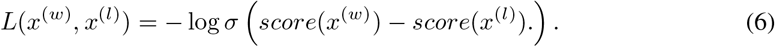

We finetuned our pretrained DNN with cross validation using the Adam optimizer and learning rate 10^*−*5^ for 15 epochs. Using cross validation on 8 random splits of 32 training examples gave an average held out Spearman correlation of *r* = 0.54. We then finetuned the pretrained model on the full dataset of 60 examples to get the final prediction model.

#### 4.1.4 VHH-feature NGS-pretrained DNN scores

We also had a version of NGS-pretrained DNN scoring system that accounted for the full variable heavy sequence, rather than just the concatenation of CDR sequences. For this scoring system, we applied very similar methods to the semi-supervised scoring system, except for the following changes: 1. For pretraining, we randomly combined the CDR sequences given by each *x*_*synth*_ with the remainder (framework) of the variable heavy chain from one of the 60 lead antibodies at random. 2. For finetuning, we used the full variable heavy chain sequences of the lead antibodies as inputs instead of just the concatenated CDR sequences. Like with our CDR-feature DNN scores, we used ESM-1v to map sequences to a vector representation, applying pretraining to the pseudo labeled sequences, and applying finetuning to the SPR-labeled antibody leads. For both pretraining and finetuning, we used 10 epochs with the Adam optimizer and learning rate 10^*−*4^. Finetuning the pretrained model with cross validation on 8 random splits of 32 training examples gave an average held out Spearman correlation of *r* = 0.52. The final model was finetuned to all 60 antibody leads.

#### 4.1.5 Final ensemble scores

The final scoring system used a weighted linear combination of the 4 scores. We determined the weighting by examining the variances of each score across the lead antibodies and single mutants of the seed antibodies. The variance approximates how much influence a score will have on the final scores, and variances can be changed by increasing or decreasing weighting. We decided to weight the CDR generative LM score the highest (based on final variance), the VHH-feature regression scores the second highest, and the two DNN scores roughly equivalent. We give the weightings and standard deviations across sequences of each score after weighting in Table 2.

**Table 2:** Weighting hyper-parameter of different scoring models used in final score and standard deviations after applying weighting. The CDR LM scores had the highest standard deviation after multiplying by weight for both single mutants of antibody seeds and across the 60 antibody leads. These standard deviations approximate how much influence a score will have on the final weighted score of a sequence–as scores that vary more will influence the final ranking more.

| Scoring model | Weight | Single mutant Std. Dev. | Antibody lead Std. Dev. |
| --- | --- | --- | --- |
| CDR generative LM | 5 | 1.62 | 2.49 |
| CDR-feature NGS-pretrained DNN | 0.6 | 0.53 | 1.07 |
| VHH-feature NGS-pretrained DNN | 0.4 | 0.39 | 1.15 |
| VHH-feature linear regression | 5 | 1.06 | 1.85 |

### 4.2 Generating antibody designs

We designed a set of 96 antibodies for laboratory testing. We used 2 different strategies for generating designs. The first involved generating between 1 to 5 mutations from top hits from the original antibody leads. The second strategy involved designing new re-combinations of the CDR sequences from the original antibody leads.

#### 4.2.1 Designing antibodies with mutational trajectories

We wanted to sample from constrained sets of *x* that correspond to specific numbers of mutations within the CDRs. We define *X*_*n*_(*x*^*wt*^) as the set of sequences that are exactly *n* mutations away from a seed protein *x*^*wt*^.

We assume that we have a scoring function *Score*(*x*) that is going to guide the sampling procedure. We would like to be able to sample from constrained sets of *x* that correspond to specific numbers of mutations. We define *M*_*n*_(*x*_*wt*_) as the set of sequences that are exactly *n* mutations away from a template protein *x*^*wt*^. An energy function based on *Score*(*x*) could be used to sample sequences *n* mutations away, given by

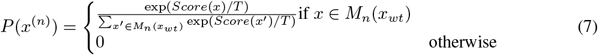

Our goal was to sample a large and diverse set of sequences that have high scores to select the final candidates from. Procedures for sampling directly from the energy function such Markov Chain Monte Carlo would work well for this with enough samples, but also result in samples that are locally correlated. We used a slightly different strategy to sample mutational trajectories to give independently generated samples that are *n* mutations away from a starting protein.

We start by sampling from Equation 7 for *n* = 1, which is tractable because it only includes single mutations (and only in CDRs). We then sample from the single mutants of (*x*^(1)^) in the same way, but exclude mutations to the position that was mutated in the first iteration. This ensures all sampled proteins will be in the set *M*_2_(*x*_*wt*_), which contains only double mutants of the original wildtype. Repeating this procedure iteratively *n* times results in n-mutant sequences. The algorithm for this process is given below.
define *M*_*n*_(*x*) as a function that returns the set of all the *n* mutant sequences of *x x*(0) *←x*_wt_

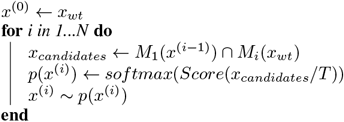

We generated sequences using 5 different antibody leads from the original library as seed sequences *x*_*wt*_. For each seed sequence, we generated 256 each of 1,2,3,4,and 5 mutant sequences from each of the 5 seed sequences using the mutational trajectory procedure.

We selectrd 15 antibodies for synthesis from each seed sequence, and chose to include 3 each of 1,2,3,4,and 5 mutant sequences in the final set. Our goal was to select a diverse set of candidates that have high scores under our scoring function *Score*(*x*). We start with a pool of candidates *X*_*pool*_ that were generated in the previous step.

For a given seed sequence, we iteratively perform sequence selection 15 times to select the highest scoring sequence that is sufficiently different from the previously selected sequences. To do this, we use our fixed ensemble scoring function *Score*(*x*), and a similarity penalty function *pen*(*x*) that is updated after each selection. For computing similarity penalties, we represent each sequence as a series of mutations from the seed sequence. We initialize two new empty dictionaries, *c*_*m*_ and *c*_*p*_, that will store the counts of each selected mutation and each selected mutation position respectively. At each iteration we use the scores and penalties to select one sequence to be moved from *X*_*pool*_ to *X*_*final*_, which is the set of sequences that will be synthesized.

We apply the following steps at each iteration:

1. Find *argmaxx*(*score*(*x*) *−λ pen*(*x*)) and add this to *X*_*final*_, and remove this sequence from *X*_*pool*_. We used *λ* = 1 to weight the penalties with the scores.
2. Get all the mutations in the previously selected sequence. Each mutation is represented as a sequence position and a new amino acid for that position. Add these mutations to the *c*_*m*_ dictionary (or increase their stored counts by 1 if they are already in the dictionary).
3. Get all the mutation positions in the most recently added sequence to *x*_*final*_. Add these mutations to the *c*_*p*_ dictionary (or increase their stored counts by 1 if they are already in the dictionary).
4. Update the similarity penalties *pen*(*x*) for the remaining sequences. For a given sequence *x*, the similarity penalty has a position component and a mutation component. We define the mutations in *x* as *x*_*m*_.

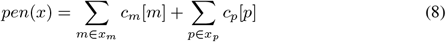

5. Check if *x*_*final*_ contains 3 n-mutant sequences from the seed sequence for all *n* from 1-5. If we have selected 3 n-mutant sequences, remove all n-mutant sequences from *X*_*pool*_.

#### 4.2.2 Designing antibodies by recombining CDR sequences

We also tested 21 sequences that were novel combinations of CDRs present in the antibody leads. We consider all possible combinations of CDR sequences using spans with non-zero frequencies in 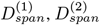, and 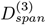. This yielded a set of 53*k* CDR-only sequences to chose from. We com-puted CDR generative LM scores and CDR-feature NGS-pretrained DNN scores only (weighting them using their corresponding weights in Table 2), since our other 2 scores rely on having the full sequence. Like with the mutational trajectories, we selected sequences one-by-one from the pool of candidates *X*_*pool*_, adding penalties for being similar to previously selected sequences. We initialized an empty dictionary *c*_*cdr*_ to store the counts of each unique CDR sequence in selected sequences. We apply the following steps at each iteration:

1. Find *argmaxx*(*score*(*x*) *−λ pen*(*x*)) and add this to *X*_*final*_, and remove this sequence from *X*_*pool*_. We used *λ* = 0.1 to weight the penalties with the scores.
2. Get all the CDR sequences of the previously selected sequence. Add the 3 CDR sequences in this sequence to the *c*_*cdr*_ dictionary (or increase their stored counts by 1 if they are already in the dictionary).
3. Update the similarity penalties *pen*(*x*) for the remaining sequences. We define *x*_*m*_ as the 3 CDR sequences in *x*.

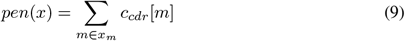

### 4.3 Collecting laboratory data

We describe methods for NGS selection and screening that were used for the anti-CD40L campaign to determine the set of antibody leads. We then describe Reformatting, expression, and purification of the antibodies to synthesize them outside of phage. Lastly we describe our SPR affinity measurements. Our methods closely follow a previous nanobody library design protocol (Yuan et al., 2022).

#### 4.3.1 Panning and screening strategy for NGS selection

Phage particles were blocked with phosphate-buffered saline (PBS) with 5% BSA and depleted for nonspecific binders on M-280 streptavidin coated magnetic beads (Thermo Fisher Scientific). Biotinylated Human CD40L (Acro CDL-H52Db) was mixed with M-280 beads (100 nM per 1 mg bead), washed with PBS/0.5% Tween to remove unbound protein and used as panning target for four rounds of panning. Phage supernatant depleted of nonspecific binders were transferred to bead mixture containing bound biotinylated CD40L and allowed to bind for 1 hour at RT to select for binders with gentle nutation. Following incubation, beads were washed several times with PBS/0.5% Tween to remove non-binding clones. Remaining bound phage were eluted trypsin in PBS buffer for 30 minutes at 37°C. The output supernatant enriched in binding clones was amplified in TG1 E. coli cells to use as input phage for the next round of selection, with each round increasing the wash cycles and lowering the total amount of antigen present.

The eluted phage pool resulting from four rounds of panning was subjected to NGS sequencing using the Illumina MiSeq platform. Assembled reads from the sequencer underwent clustering and enrichment ranking, with subsequent selection of the top 100 clusters for downstream antibody expression, purification, and characterization.

Bacterial colonies containing the phagemid display vector were isolated on 2YT agar plates with 100 *μg/ml* carbenicillin and single colonies were picked using QPix 420 (Molecular Devices) into 384-well plates containing 2YT with M13KO7 helper phage (Antibody Design Labs) to express phage for use in ELISAs. Phage ELISAs were conducted using Nunc 384-well plates (Thermo) with passively absorbed human CD40L. Anti-M13 antibody conjugated to horseradish peroxidase (HRP) (Sino Biological 11973-MM05T-H) was used to detect the presence of bound phage following the addition of 3,3’,5,5’-tetramethylbenzidine substrate. Clones that demonstrated three-fold binding over BSA background were submitted for rolling circle amplification (RCA) and Sanger sequencing to GENEWIZ using phiS4 (GCGGATAACAATTTGAATTCAAGGAGACAG) or psiR2 (CGTTAGTAAATGAATTTTCTGTATGAGG) primers to identify the VHH region.

#### 4.3.2 Reformatting, expression, and purification of antibodies

VHH single-domain antibodies were reformatted to VHH-Fc for DNA back-translation, synthesis, and cloning into mammalian expression vector pTwist CMV BG WPRE Neo utilizing the Twist Bioscience eCommerce portal. Clonal genes were delivered as purified plasmid DNA ready for transient transfection in HEK Expi293 cells (Thermo Fisher Scientific). Cultures in a volume of 1.2 mL were grown to 4 days, harvested and purified using Protein A resin (PhyNexus) on the Hamilton Microlab STAR platform into 43 mM citrate 148 mM HEPES, pH 6. CE-SDS was used to determine antibody purity and confirm molecular weight.

#### 4.3.3 SPR affinity measurements

SPR experiments were performed on a Carterra LSA SPR biosensor equipped with a HC30M chip at 25°C in HBS-TE. Antibodies were diluted to 10 μg/mL and amine-coupled to the sensor chip by EDC/NHS activation, followed by ethanolamine HCl quenching. Increasing concentrations of analyte were flowed over the sensor chip in HBS-TE with 0.5 mg/mL BSA with 5 minute association and 15 minute dissociation. CD40L was sourced from AcroBiosciences (Acro CDL-H52Db). Following each injection cycle, the surface was regenerated with 2× 30-second injections of IgG elution buffer (Thermo). Data were analyzed in Carterra’s Kinetics Tool software with 1:1 binding model.

## 5 Acknowledgements

We would like to thank Ali Madani and Richard Socher for their roles in establishing the research direction and securing resources and funding, and Hoa Giang, Maxwell Stefan, Joyce Lai, and Sejal Petal for supporting the collaboration between Salesforce and Twist.

